# Biomarker Identification by Proteomic Analysis of Vitreous Humor and Plasma in Diabetic Retinopathy

**DOI:** 10.1101/2024.05.18.594835

**Authors:** Qian Huang, Angela Banks, Rebecca Stacy, Ning Li, Yesel Kim, Lori Jennings, Nancy Finkel, Stella Yao, Anfan Wu, Amy Chen, Maen Obeidat, Cynthia Grosskreutz, S.H. Melissa Liew, Ganesh Prasanna, Hyeong Gon Yu, Joseph Loureiro, Qin Zhang

## Abstract

**Importance:** Identify detectable plasma and/or vitreous signals to potentially predict diabetic retinopathy (DR) progression for earlier disease intervention.

**Objective:** To determine the mediators and potential disease progression biomarkers of DR in vitreous humor (VH) and plasma samples using the SomaScan proteome profiling platform.

**Design:** Differential expression analysis was conducted on VH and plasma samples using the SomaScan Assay.

**Setting:** A non-interventional study conducted to collect and analyze VH and plasma samples from patients with diabetic retinopathy.

**Participants:** Samples from DR (60 nonproliferative diabetic retinopathy/NPDR, 60 proliferative diabetic retinopathy/PDR) and 60 control patients were collected.

**Main outcomes and Measures:** Differentially expressed proteins between disease and control groups were identified. Pathway enrichment analysis was conducted to identify significantly perturbed pathways in DR. Finally, a random forest model was used to identify predictive biomarkers of disease progression.

**Results:** SomaScan v3 is a pooled aptamer hybridization assay using 5080 SOMAmers to probe over 4100 proteoforms in VH and plasma samples from 3 groups (control, NPDR, and PDR). The most profound protein content change was observed in the VH samples of PDR patients, while minimal changes were measured in plasma samples, highlighting the regionality of PDR pathogenesis. Many key molecules and molecular pathways such as VEGF-A, erythropoietin, and inflammation-associated proteins implicated in DR were significantly affected in the VH of PDR patients. In addition to the classic pathways (hypoxia, immune response, mTORC1 signaling) known to be involved in PDR, novel signaling pathways, including HEME metabolism and adipogenesis, were identified in VH samples. Application of a machine learning algorithm identified a panel of plasma PDR predictive biomarkers and revealed SCARA5 as the top one based on the largest average Gini decrease in the model.

**Conclusion:** Our study identified profound alteration of protein expression and molecular pathways in the VH of PDR patients, supporting the key role of local pathogenic changes in DR progression compared to systemic factors. Although the systemic changes related to DR were small, a few disease progression predictive candidate biomarkers (SCARA5, PTK7, FAM3Band FAM3D) were identified, prompting further investigation.

**Key Points:** **Question:** Are plasma/ vitreous humor (VH) proteins predictive of diabetic retinopathy (DR) progression?

**Findings:** This study identifies substantial protein changes in the VH of proliferative diabetic retinopathy (PDR) patients, while early nonproliferative DR (NPDR) patients show minimal change. We identify multiple proteins linked to angiogenesis, inflammation, immune cells (microglia/macrophage/neutrophil), and leukostasis associated with PDR and reveal a potential plasma panel of disease progression (from NPDR to PDR) biomarkers (*SCARA5, PTK7, FAM3B, FAM3D*).

**Meaning:** Identified disease progression predictive biomarkers permits potential development of prognostic tools to identify individuals most at risk for PDR progression and offering reduced disease burden by earlier intervention.

## Introduction

Diabetic retinopathy (DR) is a common and specific microvascular complication of diabetes. This pathological condition can cause severe damage to the retina^1^ leading to vision loss and blindness among working age adults in developed countries.^2^ DR can be further classified into 2 types: nonproliferative diabetic retinopathy (NPDR) and proliferative diabetic retinopathy (PDR). Classification of NPDR is based on clinical findings manifested by visible vascular features, including microaneurysms, retinal hemorrhages, intraretinal microvascular abnormalities, and venous caliber changes, while PDR is characterized by the hallmark feature of pathologic preretinal neovascularization.^3^ Many risk factors (e.g., hyperglycemia, hypertension) are believed to initiate a cascade of biochemical and physiological changes that ultimately lead to microvascular damage and retinal dysfunction.^4^ Despite recent advances in the treatment of DR, there remains a large unmet clinical need to prevent or reverse vision loss by treating earlier stages of the disease. Therefore, novel biomarkers are needed to identify individuals most at risk for PDR progression and intervene early, thus limiting vision loss and reducing the cost associated with managing the more advanced disease. The purpose of this study was to identify predictive biomarkers in vitreous humor (VH) or plasma for DR progression.

The physiologic and pathophysiologic conditions of the retina are most likely reflected in the protein composition of the VH, due to their close anatomical and biological relationship, and can be sampled as part of routine surgical procedures.^5^ Large-scale protein studies using proteomic techniques have proved very effective in generating novel knowledge about the complex pathological mechanisms underlying DR.^6,7^

In this study, we have carried out a proteomics profiling of the VH alongside plasma from healthy and DR patients by SomaScan analysis. We examined the protein profiles of human VH from patients with PDR or NPDR. There remains a major unmet medical need to develop biomarkers that can be used in a routine clinical setting to identify individuals with DR who will develop PDR.

The comprehensive SomaScan analysis provides the means to identify key nodes or pathways that are disrupted in the retina of diabetic patients and to detect plasma and vitreous signals which can predict DR progression.

## Methods

### Subject Enrollment and Sample Collection

This non interventional study was approved by the Novartis BioMedical Research (NBR) in collaboration with Seoul National University Hospital in Korea. Signed informed consent was obtained from every participant before being included into the study. VH and plasma samples were collected from patients who underwent a standard of care vitrectomy. A total of 180 subjects, 60 control (nondiabetic controls including but not restricted to patients with epiretinal membrane, retinal detachment, vitreous hemorrhage, and idiopathic macular hole), 60 NPDR, and 60 PDR were enrolled. The inclusion criteria were as follows: patient age between 18 and 85 years, a need to have vitrectomy as part of their standard of care for conditions including but not limited to DR, retinal detachment, macular hole, vitreous hemorrhage, epiretinal membrane, or diabetes-related complications. The exclusion criteria were as follows: history of neovascular glaucoma or inflammatory glaucoma (e.g., uveitic, Posner-Schlossman syndrome) in the study eye or a history of glaucoma surgery in the study eye. However, patients with primary open-angle glaucoma with no history of surgical treatment were included.

### Sample Preparation

The plasma and VH samples were collected and stored at −80°C. VH were added with 1:100 Halt Protease and phosphatase inhibitor cocktail (Thermo Scientific, #1861280), 1:200 PMSF (Cell Signaling #8553S), and 1:500 EDTA (Thermo Scientific, #R1021) according to the weight, then subjected to centrifugation at 10 000 rpm 4°C for 2 minutes. An aliquot of the plasma and VH samples was sent to SomaLogic for analysis.

### SomaLogic Analysis

To generate a proteomic dataset of the 180 patients we used the SomaScan assay to measure ∼4000 proteins with high affinity and specificity. The SomaScan v3 used measures 5080 SOMAmers per sample. Two batches of plasma and vitreous samples were sent to SomaLogic Inc. (Boulder, Colorado, United States) for protein profiling.^8^

### Data Analysis

SOMAmer relative fluorescent units matrices for all samples were analyzed internally. Quality control and sample outlier detection were conducted using the arrayQualityMetrics R package,^9^ and by running principal component analysis (PCA) with custom R scripts. Batch effects were evaluated using PCA and mitigated using the ComBat method.^10^ Batch-corrected, relative fluorescent units were log2-transformed and normalized by smooth quantile normalization.^11^ We applied the linear models for microarray data (limma) R package to determine differentially expressed proteins between disease groups, while adjusting for age- and sex-related effects.^12^ Significantly differentially expressed proteins were defined as proteins which had an adjusted *P* value ≤ 0.05 and log2 fold change (FC) ≥ 0.5. Pathway level changes were investigated with gene set enrichment analysis with the fgsea R package.^13^ To identify biomarkers predictive of disease progression, we built a random forest model using 75% of the plasma sample data, focusing on proteins that are differentially expressed in PDR. The accuracy of the model was assessed using the remaining 25% of the sample data. Top predictive biomarkers were defined as proteins with the largest average Gini decrease in the model.

## Results

### Sample Demographics

We collected and analyzed paired VH and plasma samples from 180 patients. The age of patients in the NPDR group was significantly older than the control and PDR groups. There was no significant age difference between the PDR and the control group (Supplementary eFigure 1A). The gender distribution is shown in Supplementary eFigure 1B.

**Figure 1:**
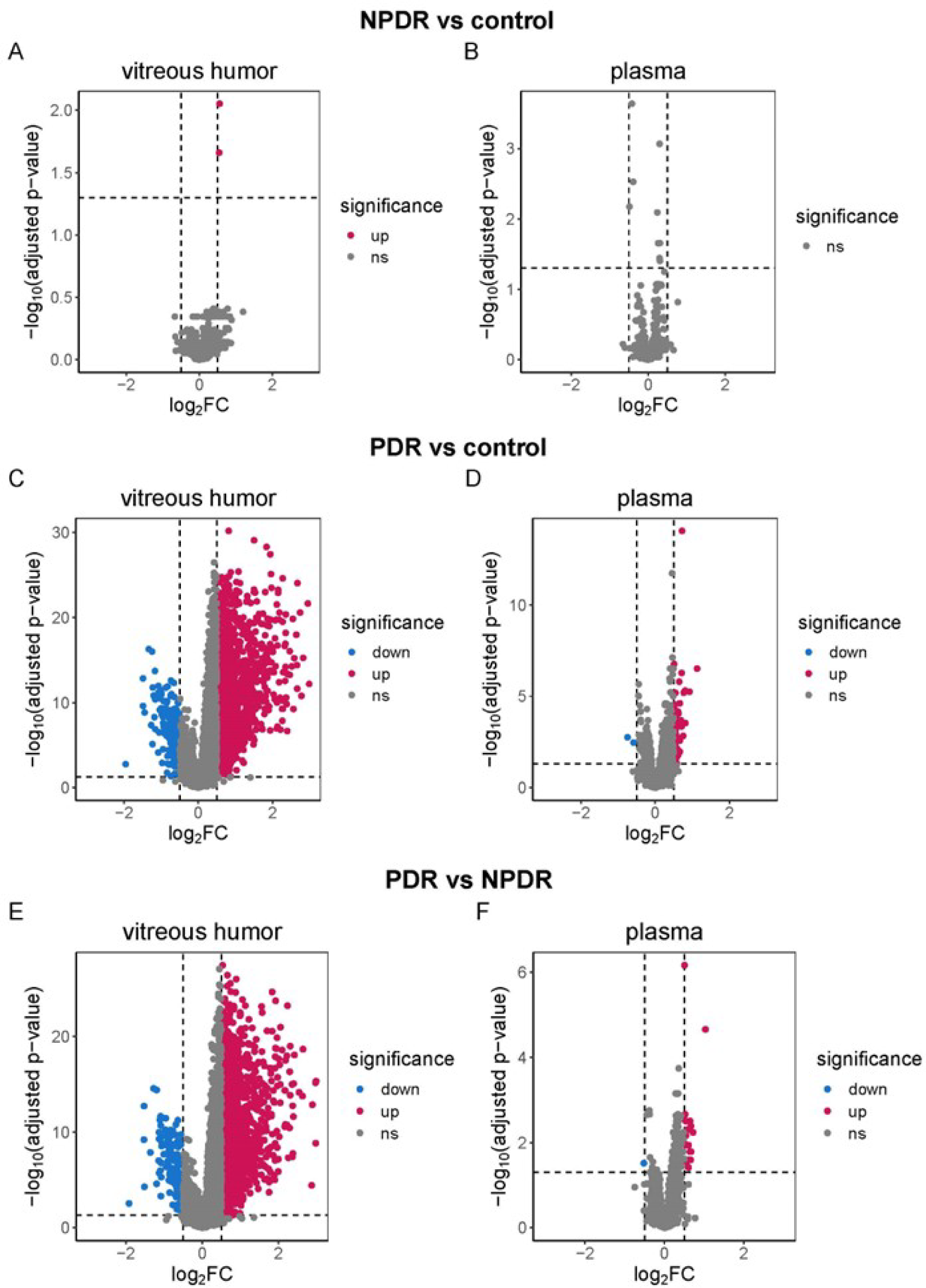
Differential Expression Analysis Revealed the Greatest Protein Abundance Change in the VH of PDR Patients. Differentially expressed proteins shown in a volcano plot between-group comparison between NPDR and nondiabetic control in VH (Figure 1A) and plasma (Figure 1B); PDR vs nondiabetic control in VH (Figure 1C) and plasma (Figure 1D); and PDR vs NPDR in VH (Figure 1E) and plasma (Figure 1F). Red dots show significantly upregulated proteins and blue dots indicate the significantly downregulated proteins based on log2FC ≥ 0.5, with multiple comparison correction using false discovery rate applied at a threshold of *P* value ≤ 0.05). FC, fold change; FDR, false discovery rate; NPDR, nonproliferative diabetic retinopathy; ns, not significant; PDR, proliferative diabetic retinopathy; VH, vitreous humor.

### Differential Expression Analysis of PDR, NPDR, and Nondiabetic Controls Shows Substantial Changes at the PDR Stage, While Minimal Changes at the NPDR Stage

To investigate changes in protein abundance between nondiabetic controls and NPDR patients in the VH and plasma, we performed differential expression analysis using the limma method.^12^ Interestingly, we found few differential proteins between NPDR patients and nondiabetic controls in the VH or plasma (log2FC ≥ 0.5, FDR-corrected *P* value ≤ 0.05; Figure 1A & 1B), indicating that only subtle protein abundance changes occur in the earlier stages of DR. However, there was a significant amount of protein expression change in the PDR VH (Figure 1C). A handful of protein expression changes were also in PDR plasma (Figure 1D).

The only protein significantly upregulated in the NPDR VH was MMP10; no changes of MMP10 in NPDR plasma were observed. In NPDR plasma, several proteins were altered with a significant *P* value, but the log2FC values were ≤ 0.5. These upregulated proteins included PTK7, NFASC, IGSF3, IL1R1, SEMA6B, SEMA6A, and downregulated proteins were CXCL12, EPYC, and CCL22.

### PDR Changes Occur Locally in the Vitreous Humor

Next, we investigated changes in protein abundance between PDR patients and nondiabetic controls, as well as NPDR. In the VH of PDR patients, we found robust protein abundance changes, with 163 significantly downregulated proteins and 1332 significantly upregulated proteins compared to nondiabetic controls (log2FC ≥ 0.5, FDR-corrected *P* value ≤ 0.05; Figure 1C, Supplementary Excel file). Similar findings were obtained when we compared PDR with NPDR, with 170 significantly downregulated proteins and 1241 significantly upregulated proteins in PDR VH (log2FC ≥ 0.5, FDR-corrected *P* value ≤ 0.05; Figure 1E, Supplementary Excel file). In contrast, the plasma of PDR patients showed relatively minimal protein changes, with only 2 significantly downregulated proteins (FTH1 and FAM189A2) and 45 significantly upregulated proteins compared to nondiabetic controls (log2FC ≥ 0.5, FDR-corrected *P* value ≤ 0.05, Figure 1D, Supplementary Excel file). Only 1 significantly downregulated protein (ACY1) and 20 significantly upregulated proteins were identified in PDR plasma compared to NPDR (Figure 1F, Supplementary Excel file). The majority of these upregulated proteins (SCARA5, CSF1, APOF, PXDN, TNFRSF1A, NBL1, REN, ROR2, TFF3, CHGA, FAM3D, IGFBP1, IGFBP2, LCN2, DEFA5, RNASE1, HAVCR1, TAGLN) are increased in the PDR plasma compared to both nondiabetic controls and NPDR. These results indicate that DR likely leads to local changes in the eye, rather than systemic changes.

### Key Molecules and Molecular Pathways Implicated in DR Were Significantly Affected in the VH of PDR Patients

Diabetes reportedly affects the entire neurovascular of the retina, inducing gradual angiogenesis, inflammation, and breakdown of the retinal barrier.^14^ The consequences of these molecular events include compromised capillary function, infiltrating immune cells, tissue edema, hypoxia, and neurodegeneration. We aimed to explore each one of these earlier molecular events in detail to determine whether key proteins involved in these pathways are significantly perturbed in DR by analyzing VH.

### Vascular Endothelial Growth Factors

Vascular endothelial growth factors (VEGFs) are a family of secreted polypeptides with a highly conserved receptor-binding cystine-knot structure comprise five members: VEGF-A, placental growth factor (PGF), VEGF-B, VEGF-C, and VEGF-D.^15^ VEGF-A plays a central role in mediating microvascular and macrovascular pathology in diabetes. Diabetes leads to overexpression of angiogenic factor VEGF-A due to the hyperglycemic environment and further upregulation by tissue hypoxia.^16^ Anti-VEGF-A therapies have proven to be a successful treatment for PDR.^17^ As a proof of concept, we investigated how VEGF family members were affected in our PDR patients VH. As expected, SOMAmers for VEGF-A were significantly upregulated in the PDR group compared to the nondiabetic control group or the NPDR group (Figure 2A), but not VEGF-B and VEGF-C SOMAmers, which are weakly involved in angiogenesis.^18,19^

**Figure 2.**
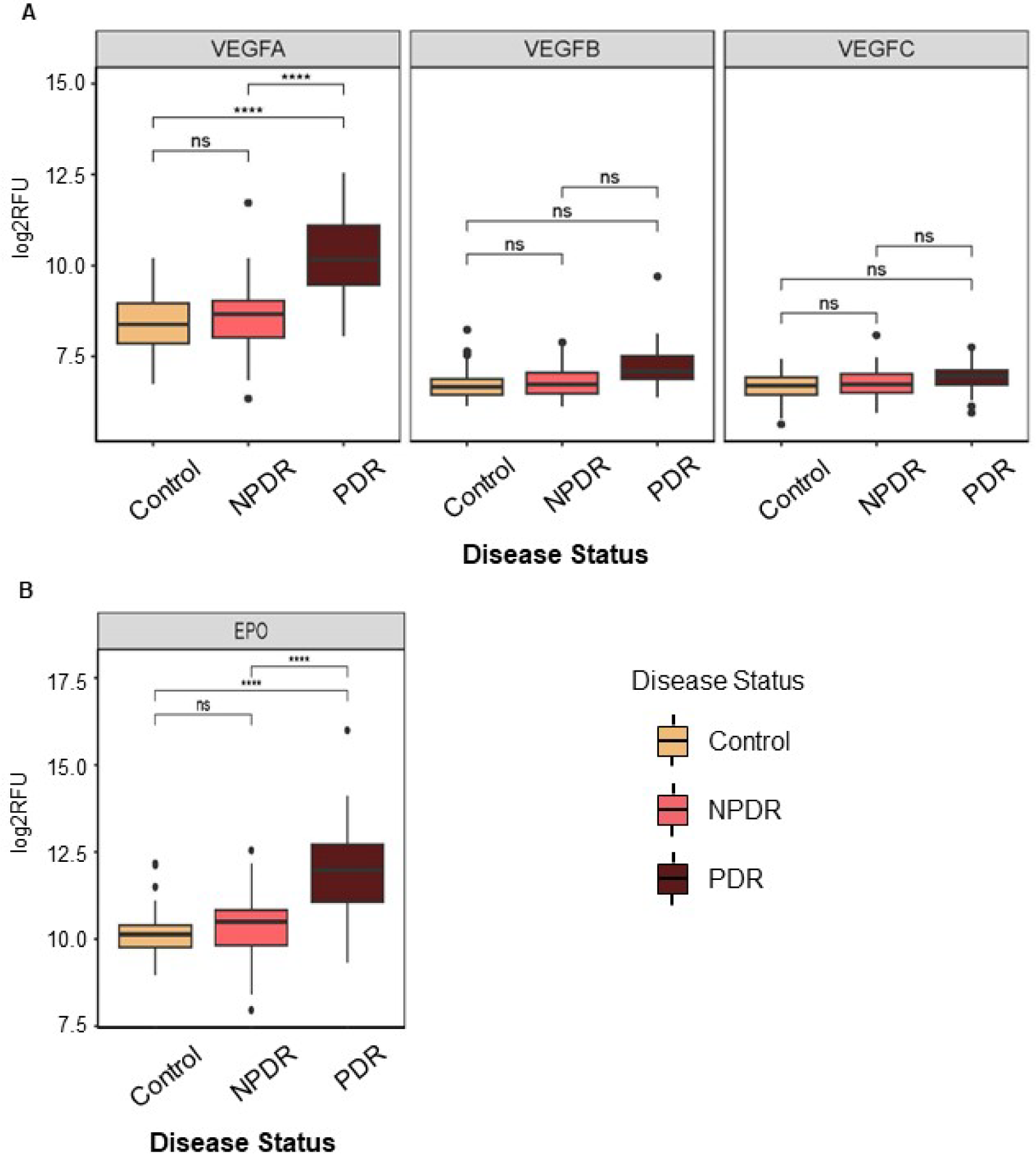
Vascular Endothelial Growth Factors (VEGF) and Erythropoietin (EPO) in the VH of PDR Patients. Angiogenesis-associated molecules VEGF-A and EPO were significantly upregulated in the vitreous humor (VH) of proliferative (PDR) patients (Figure 2A and Figure 2B) but not VEGF-B and VEGF-C molecules. ****, correction for multiple comparisons using FDR correction was applied at threshold of p≤1e-04. EPO, Erythropoietin; FDR, false discovery rate; NPDR, nonproliferative diabetic retinopathy; ns, not significant; PDR, proliferative diabetic retinopathy; RFU, relative fluorescent unit; VEGF, vascular endothelial growth factors.

### Erythropoietin (EPO)

Interestingly, other proteins with angiogenic activity were also upregulated in PDR VH, including EPO (Figure 2B). EPO, a 30.4 kDa glycoprotein hormone, is the principal regulator of erythropoiesis, by inhibiting apoptosis and stimulating the proliferation and differentiation of erythroid precursor cells.^20^ EPO is expressed in the human retina, and it is highly elevated in the VH of patients with diabetic macular edema.^21^ EPO could enhance the effects of VEGF-A, thus contributing to neovascularization and worsening PDR.^22^

### Inflammation

It has been well established that patients with DR show increased glycolytic metabolites in the eye, thereby triggering an inflammatory response.^23^ Our data support the role of inflammation in DR disease. Multiple cytokines, chemokines, and adhesion molecules were upregulated in the VH of our PDR patients (Figure 3). These proteins included members of the IL-1 family and the IL-17 family, as well as the cadherin family. Further, we noted several monocyte and macrophage markers, as well as neutrophil markers, upregulated in the PDR VH. These observations indicate that PDR induces the infiltration of immune cells and an inflammatory response.

**Figure 3.**
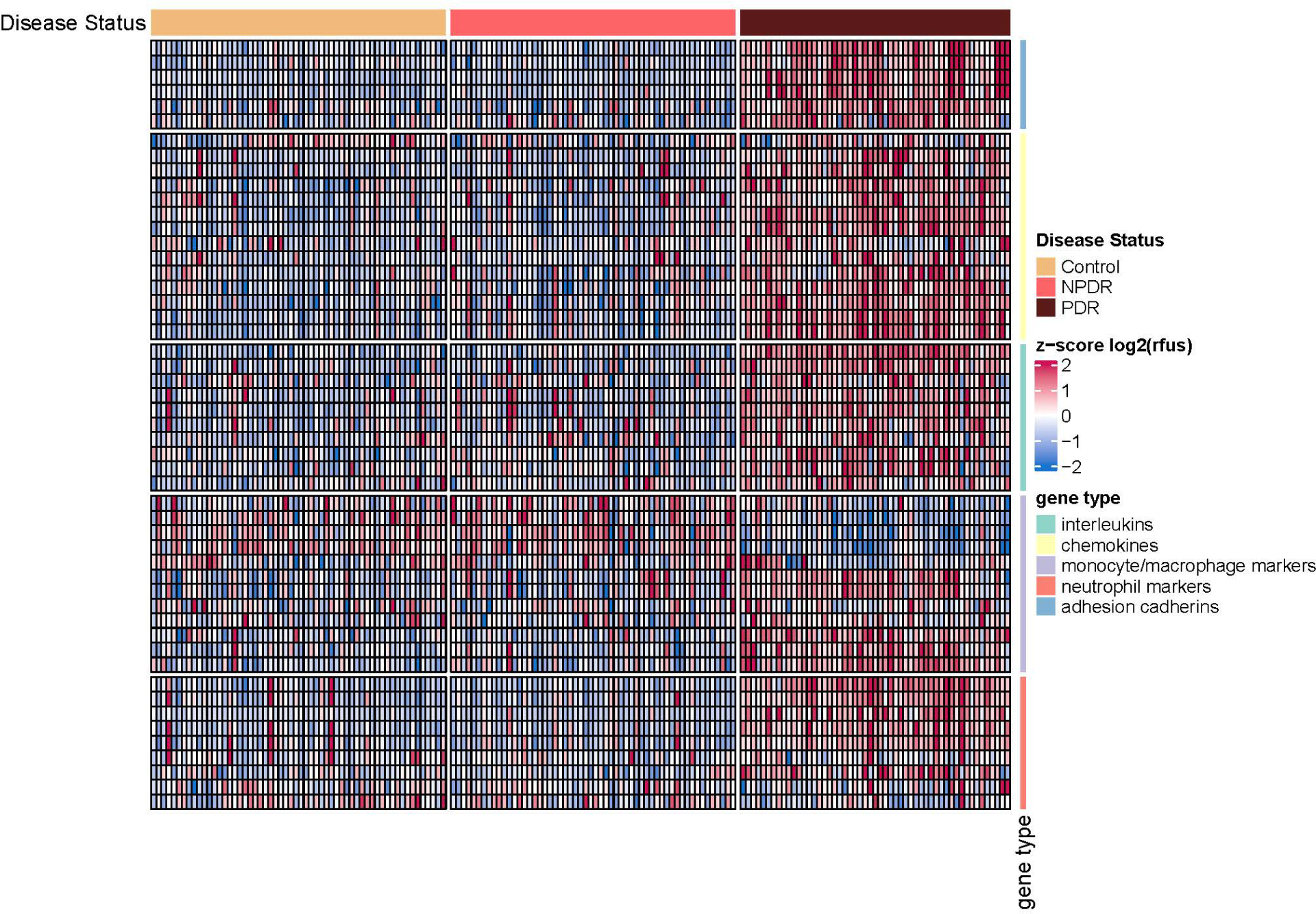
Inflammation in the VH of PDR Patients. Key molecules involved in inflammation, such as cytokines/chemokines, monocyte/macrophage/neutrophil markers, and adhesion molecules, were significantly affected in the VH of PDR patients as shown by the consistent pattern of upregulation (higher z-scores as indicated by red sections identified in Figure 3) evident across gene types (interleukins, chemokines, monocyte/macrophage markers, neutrophil makers and adhesion cadherins). VH, vitreous humor; NPDR, nonproliferative diabetic retinopathy; PDR, proliferative diabetic retinopathy.

### Pathway Analysis

Many PDR-associated pathways and events, such as hypoxia, complement, inflammatory response, coagulation, mTORC1 signaling, and epithelial mesenchymal transition (EMT), were significantly upregulated in PDR VH (Table 1). We have also identified a few novel pathways perturbed in PDR, such as HEME metabolism, and adipogenesis. In PDR plasma sample analysis, we identified known PDR-associated pathways including angiogenesis and inflammation.

**Table 1:**
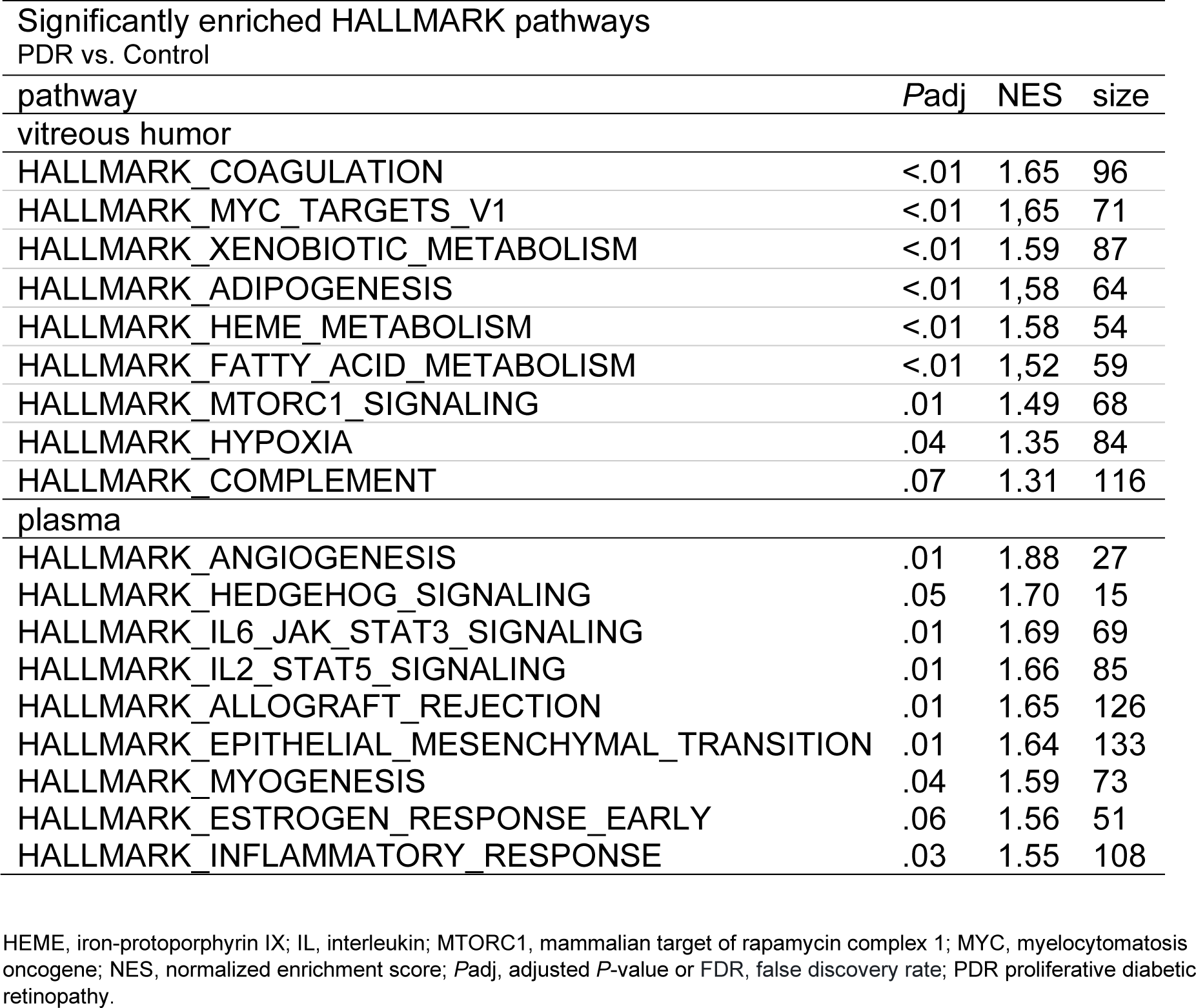
Pathways Upregulated in PDR Patients Based on fgsea.

### Discovery of Potential Disease Progression Plasma Biomarkers

The advanced high-throughput proteomic SomaLogic study enables us to examine DR plasma samples for potential DR biomarkers, particularly the disease progression predictive biomarkers. In our study, by building a machine learning algorithm with random forest using 75% of our plasma sample data, we were able to identify SCARA5 (scavenger receptor class A, member 5), PTK7 (protein tyrosine kinase 7), FAM3B (family with sequence similarity 3 member B, also known as PANDER, Pancreatic-derived factor),and FAM3D (family with sequence similarity 3 member D) as the top biomarkers (Figure 4) for classifying control, NPDR, and PDR patients based on their average decreased Gini scores (4.20, 3.50, 2.42, and 1.43 respectively). These proteins are promising candidate biomarkers for predicting DR disease progression.

**Figure 4:**
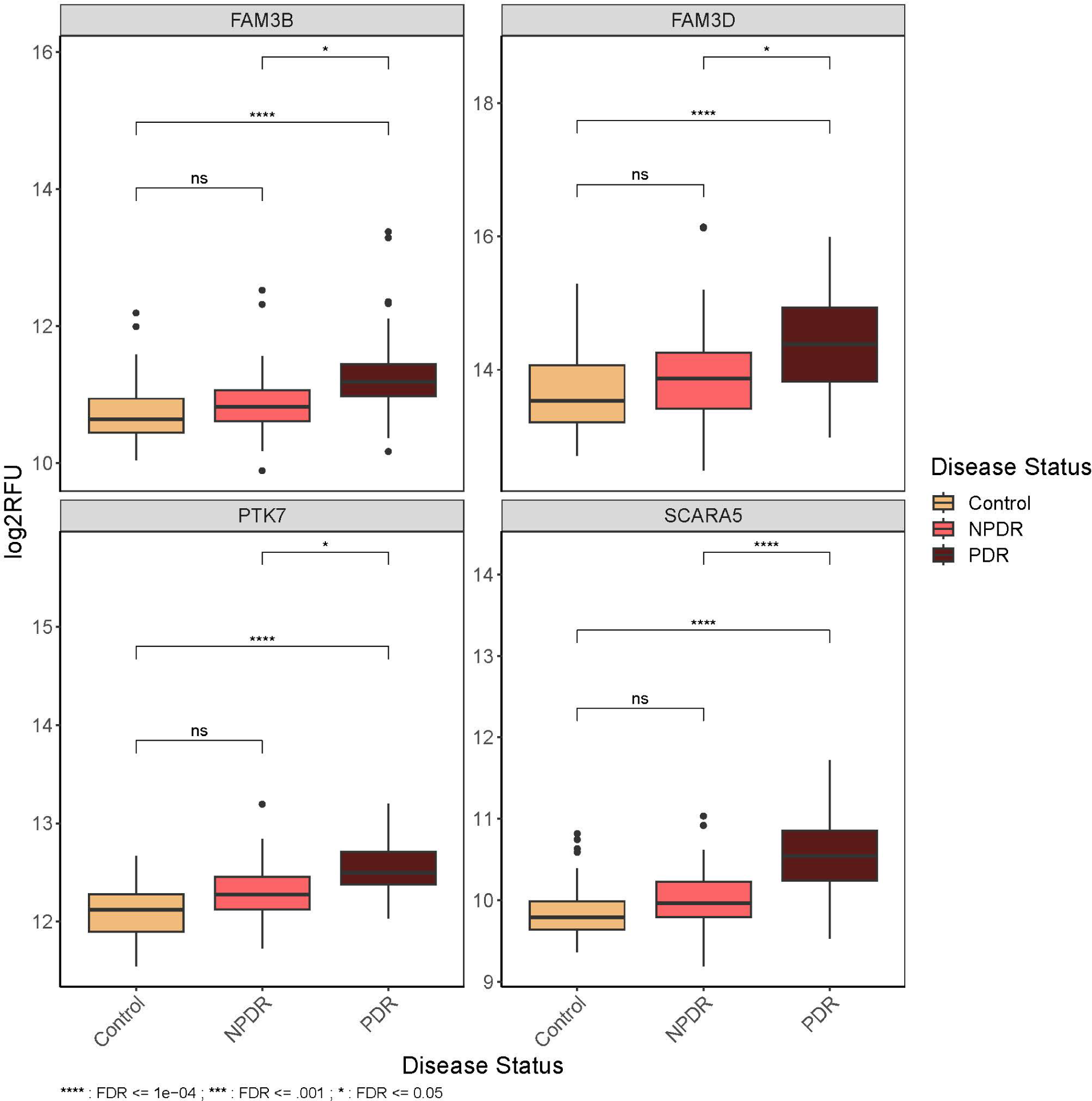
Predictive PDR Plasma Biomarkers. SCARA5 involving iron ion homeostasis was significantly upregulated in plasma of proliferative DR patients and identified as the top PDR predictive biomarker by a machine learning algorithm. Followed by PTK7, FAM3B and FAM3D. ****, correction for multiple comparisons using FDR correction was applied at threshold of p≤1e-04, ***, FDR <=0.001, *, FDR <=0.05. FAM3B, family with sequence similarity 3 member B, also known as PANDER, Pancreatic-derived factor; FAM3D, family with sequence similarity 3 member D; FDR, false discovery rate; NPDR, nonproliferative diabetic retinopathy; ns, not significant; PDR, proliferative diabetic retinopathy; PTK7, protein tyrosine kinase 7; RFU, relative fluorescent unit; SCARA5, Scavenger Receptor Class A, member 5.

## Discussion

In this study, we have analyzed proteomic changes in VH and plasma samples from nondiabetic control, NPDR, and PDR patients. Interestingly, we found minimal differential proteins between NPDR patients and nondiabetic controls, both in plasma and VH. MMP10 is the only protein significantly upregulated in the VH of NPDR patients; it has been reported that MMP10 is involved in the development of microvascular complications in diabetes, a higher MMP10 level is associated with a higher risk of retinopathy.^24^ When DR progressed to a more advanced stage (PDR), we found substantial proteomic changes. There were more than 160 proteins downregulated and 1300 proteins upregulated in the VH of PDR and 2 proteins downregulated and 45 proteins upregulated in the plasma of PDR compared to nondiabetic controls. These data demonstrate substantial proteomic changes in ocular or systemic biofluids only observed in PDR, a more advanced stage of DR, but not in the early stage (NPDR). These data also suggest that DR leads to local changes in the eye, rather than systemic changes, which provides the rationale for local treatment in a routine clinical setting.^25^

Proteomic studies examining the ocular and systemic biofluids of diabetic patients have made an important contribution to the advancement of our understanding of DR pathogenesis.^23^ High-throughput protein analysis techniques such as SomaScan have great potential for biomarker discovery (diagnostic and prognostic) and therapeutic monitoring, as well as identification of novel therapeutic targets. Intravitreal injection of anti-VEGF is considered as first-line pharmacological therapy.^25^ In our study, we demonstrated significant upregulation of VEGF-A in PDR patients’ vitreous, but not in plasma. We also found significant upregulation of EPO in PDR patients’ VH. Angiogenesis is caused by the activation of HIF-1*α* under hypoxic conditions, promoting expression of VEGF-A transcript. At the same time, regulation of EPO is also under the control of HIF-1*α*.^26^

In earlier stages of NPDR, damage to the retinal microvasculature occurs as a consequence of chronic hyperglycemia and manifests as microaneurysms, intraretinal hemorrhages, and the development of diabetic macular edema. Without appropriate treatment, the resulting retinal ischemia promotes neovascularization characteristic of PDR, leading to vitreous hemorrhage, retinal detachment, neovascular glaucoma, and severe vision loss.^27^ The main cause of NPDR to PDR progression is represented by extensive retinal ischemia, which promotes vitreoretinal neovascularization. The formation of new blood vessels is also supported by activation of retinal immune cells: microglia, monocyte-derived tissue macrophage.^28^ Our data have shown increased local expression of proinflammatory cytokines/chemokines, microglia/macrophage-associated proteins, and neutrophil-associated markers in the VH of PDR (Figure 3), which suggest that all these inflammatory responses may contribute to neovascularization in the retina during DR. The combination of increased growth factors, like VEGF-A, increased cytokines and inflammation play a causal role in breaking down the blood-retinal barrier, ultimately leading to vision loss in DR patients.^29^

Leukocyte-endothelium interactions mediated by adhesion molecules (eg, ICAM1 and SELL, which were all upregulated in PDR VH; Figure 3) have been implicated in leukostasis during diabetes. The upregulation of adhesion molecules promotes the binding of monocytes and leukocytes to retinal endothelial cells, which may cause vascular cell death.^30^ Our SomaScan data provide molecular evidence of retinal barrier breakdown in the VH of PDR patients.

Pathway analysis in our dataset reveals many upregulated pathways in PDR VH, including several known pathways:^31^ hypoxia, angiogenesis, complement, glycolysis, cellular response to TNF, chemotaxis, regulation of immune response, leukocyte migration, cellular response to IL-1, blood coagulation, platelet degranulation, fatty acid metabolism, epithelial mesenchymal transition, mTORC1 signaling, MAPK cascade, and KRAS. Some novel pathways, including HEME metabolism, adipogenesis, and protein folding, were identified in the VH of PDR patients. These findings indicate that diverse biological mechanisms are involved in PDR pathogenesis.

The pathogenesis and diagnosis of DR are very complex and not fully understood. Prognostic biomarkers to predict disease progression could lead to better diagnosis and early treatment. Therefore, in the current study using a machine learning approach, we aimed to identify potential biomarkers that indicate the progression of DR and can be used in early diagnosis and the management of detrimental late-stage PDR. Multiple risk factors and pathways impact the development and progression of DR (Figure 3, Table 1); hence, a multiple component biomarker panel may be more useful than separate biomarkers for evaluation using plasma samples. We applied a machine learning model using random forest and identified a panel of disease progression (from NPDR to PDR) predictive biomarkers (SCARA5, PTK7, FAM3B, and FAM3D) in plasma.

SCARA5 reportedly functions as a renal receptor for ferritin during endocytosis and delivers this iron-containing ligand to specific kidney tissues.^32^ Careful control of iron availability in the retina is central to maintenance of iron homeostasis, as its imbalance is associated with oxidative stress and the progression of several retinopathies, including DR. Ferritin, known for its role in iron storage and detoxification, has also been proposed as an iron-transporter protein through its binding to SCARA5 and TIM2 membrane receptors.^33^ Serum ferritin is transported across the blood-retinal barrier into the retina through SCARA5 binding.^34^ L-Ferritin binding to SCARA5, a new iron traffic pathway, is potentially implicated in retinopathy.^34^ SCARA5 was identified as the top DR progression predictive marker and it was also upregulated in PDR VH in our study. PTK7 is a catalytically defective receptor protein tyrosine kinase and promotes angiogenesis as well as migration and invasion of endothelial cells. ^35,36^ FAM3B and FAM3D belong to a cytokine-like family composed of 4 members, referred as FAM3A, FAM3B, FAM3C and FAM3D. FAM3B plays an important role in the glucose homeostasis, and it also facilitates vascular smooth muscle cells proliferation and migration. ^37^ FAM3D, a gut secreted protein and chemoattractant for neutrophils and monocytes. ^38^ In our dataset, FAM3D is also significantly upregulated in VH of PDR patients, however not PTK7 and FAM3B. The next step is to confirm these findings by orthogonal methods such as ELISA.

In summary, the SomaScan platform has been used for proteomics-based studies in large cohorts. The current study not only provides pathophysiological insights into disease progression, but also demonstrates the associated potential predictive biomarkers for DR progression.

## Acknowledgements

Special thanks to Katie Li, Alice Jeong for coordinate clinical site, Henry Wu, Quint Medley for sample preparation support, Carlton Dsouza for clinical data management.

## Data Sharing Statement

The reader can request the raw data (anonymized) and related documents (e.g., protocol, reporting and analysis plan, clinical study report) that underlie the results reported in this article by connecting to https://www.clinicalstudydatarequest.com and signing a Data Sharing Agreement with Novartis. These will be made available to qualified external researchers, with requests reviewed and approved by an independent review panel on the basis of scientific merit.

## Author Contributions

Dr Zhang and A Banks had full access to all of the data in the study and take responsibility for the integrity of the data and the accuracy of the data analysis. Study concept and design: Qian Huang, Rebecca Stacy, Cynthia Grosskreutz, S.H. Melissa Liew.

Acquisition: Hyeong Gon Yu.

Analysis: Qin Zhang, Angela Banks, Ning Li.

Interpretation of data: Qin Zhang, Angela Banks.

Drafting of the manuscript: Qin Zhang, Angela Banks.

Critical revision of the manuscript for important intellectual content: All authors.

Statistical analysis: Angela Banks, Ning Li.

Administrative, technical, or material support: Lori Jennings, Nancy Finkel, Joseph Loureiro, Amy Chen, Maen Obeidat.

Study supervision: Rebecca Stacy, Yesel Kim, Stella Yao, Anfan Wu, S.H. Melissa Liew.

## References

1. Antonetti DA, Klein R, Gardner TW. Diabetic retinopathy. N Engl J Med. 2012;366(13):1227–1239.

2. Liu Y, Swearingen R. Diabetic Eye Screening: Knowledge and Perspectives from Providers and Patients. Curr Diab Rep. 2017;17(10):94.

3. Stitt AW, Curtis TM, Chen M, et al. The progress in understanding and treatment of diabetic retinopathy. Prog Retin Eye Res. 2016;51:156–186.

4. Cheung N, Mitchell P, Wong TY. Diabetic retinopathy. Lancet. 2010;376(9735):124–136.

5. Cehofski LJ, Honore B, Vorum H. A Review: Proteomics in Retinal Artery Occlusion, Retinal Vein Occlusion, Diabetic Retinopathy and Acquired Macular Disorders. Int J Mol Sci. 2017;18(5):907.

6. Merchant ML, Klein JB. Proteomics and diabetic retinopathy. Clin Lab Med. 2009;29(1):139–149.

7. Weber SR, Zhao Y, Gates C, et al. Proteomic analyses of vitreous in proliferative diabetic retinopathy: prior studies and future outlook. J Clin Med. 2021;10(11):2309.

8. Gold L, Ayers D, Bertino J, et al. Aptamer-based multiplexed proteomic technology for biomarker discovery. Plos one. 2010;5(12):e15004.

9. Kauffmann A, Gentleman R, Huber W. Array Quality Metrics-a bioconductor package for quality assessment of microarray data. Bioinformatics. 2009;25(3):415–416.

10. Johnson WE, Li C, Rabinovic A. Adjusting batch effects in microarray expression data using empirical Bayes methods. Biostatistics. 2007;8(1):118–127.

11. Hicks SC, Okrah K, Paulson JN, Quackenbush J, Irizarry RA, Bravo HC.. Smooth quantile normalization. Biostatistics. 2018;19(2):185–198.

12. Ritchie ME, Phipson B, Wu D, et al. limma powers differential expression analyses for RNA-sequencing and microarray studies. Nucleic Acids Res. 2015;43(7):e47.

13. Korotkevich G, Sukhov V, Budin N, Shpak B, Artyomov MN, Sergushichev A. Fast gene set enrichment analysis. bioRxiv. 2021; doi: 10.1101/06001

14. Abcouwer SF. Angiogenic Factors and Cytokines in Diabetic Retinopathy. J Clin Cell Immunol. 2013; Suppl 1(11):1–12.

15. Holmes DI, Zachary I. The vascular endothelial growth factor (VEGF) family: angiogenic factors in health and disease. Genome Biol. 2005;6(2):209.

16. Capitao M, Soares R. Angiogenesis and Inflammation Crosstalk in Diabetic Retinopathy. J Cell Biochem. 2016;117(11):2443–2453.

17. Crawford TN, Alfaro DV 3rd, Kerrison JB, Jablon EP. Diabetic retinopathy and angiogenesis. Curr Diabetes Rev. 2009;5(1):8–13.

18. Bry M, Kivela R, Leppänen VM, Alitalo K. Vascular endothelial growth factor-B in physiology and disease. Physiol Rev. 2014;94(3):779–794.

19. Bui HM, Enis D, Robciuc MR, et al. Proteolytic activation defines distinct lymphangiogenic mechanisms for VEGFC and VEGFD. J Clin Invest. 2016;126(6):2167–2180.

20. Hernández C, Simó R. Erythropoietin produced by the retina: its role in physiology and diabetic retinopathy. Endocrine. 2012;41(2):220–226.

21. Hernández C, Fonollosa A, García-Ramírez M, et al. Erythropoietin is expressed in the human retina and it is highly elevated in the vitreous fluid of patients with diabetic macular edema. Diabetes Care. 2006;29(9):2028–2033.

22. Grant MB, Boulton ME, Ljubimov AV. Erythropoietin: when liability becomes asset in neurovascular repair. J Clin Invest. 2008;118(2):467–470.

23. Youngblood H, Robinson R, Sharma A, Sharma S. Proteomic Biomarkers of Retinal Inflammation in Diabetic Retinopathy. Int J Mol Sci. 2019;20(19):4755.

24. Toni M, Hermida J, Goni MJ, et al. Matrix metalloproteinase-10 plays an active role in microvascular complications in type 1 diabetic patients. Diabetologia. 2013;56(12):2743–2752.

25. Amoaku WM, Ghanchi F, Bailey C, et al. Diabetic retinopathy and diabetic macular oedema pathways and management: UK Consensus Working Group. Eye (Lond). 2020;34(Suppl 1):1–51.

26. Maes C, Carmeliet G, Schipani E. Hypoxia-driven pathways in bone development, regeneration and disease. Nat Rev Rheumatol. 2012;8(6):358–366.

27. Moshfeghi A, Garmo V, Sheinson D, Ghanekar A, Abbass I. Five-Year Patterns of Diabetic Retinopathy Progression in US Clinical Practice. Clin Ophthalmol. 2020;14:3651–3659

28. Dejda A, Mawambo G, Cerani A, et al. Neuropilin-1 mediates myeloid cell chemoattraction and influences retinal neuroimmune cross-talk. J Clin Invest. 2014;124(11):4807–4822.

29. Forrester JV, Kuffova L, Delibegovic M. The Role of Inflammation in Diabetic Retinopathy. Front Immunol. 2020;11:583687

30. Chibber R, Ben-Mahmud BM, Chibber S, Kohner EM. Leukocytes in diabetic retinopathy. Curr Diabetes Rev. 2007;3(1):3–14.

31. Xia HQ, Yang JR, Zhang KX, et al. Molecules related to diabetic retinopathy in the vitreous and involved pathways. Int J Ophthalmol. 2022;15(7):1180–1189.

32. Li JY, Paragas N, Ned RM, et al. Scara5 is a ferritin receptor mediating non-transferrin iron delivery. Dev Cell. 2009;16:35–46.

33. Valença A, Mendes-Jorge L, Bonet A, et al. TIM2 modulates retinal iron levels and is involved in blood-retinal barrier breakdown. Exp Eye Res. 2021; 202:108292.

34. Mendes-Jorge L, Ramos D, Valença A, et al. L-ferritin binding to scara5: a new iron traffic pathway potentially implicated in retinopathy. Plos One. 2014;9(9): e106974.

35. Shin WS, Maeng YS, Jung JW, Min JK, Kwon YG, Lee ST. Soluble PTK7 inhibits tube formation, migration, and invasion of endothelial cells and angiogenesis. Biochem Biophys Res Commun. 2008;371(4):793–798. doi:10.1016/j.bbrc.2008.04.168

36. Oh SW, Shin WS, Lee ST. Anti-PTK7 Monoclonal Antibodies Inhibit Angiogenesis by Suppressing PTK7 Function. Cancers (Basel). 2022;14(18):4463. Published 2022 Sep 14. doi:10.3390/cancers14184463

37. Zhang W, Chen S, Zhang Z, Wang C, Liu C. FAM3B mediates high glucose-induced vascular smooth muscle cell proliferation and migration via inhibition of miR-322-5p. Sci Rep. 2017;7(1):2298. Published 2017 May 23. doi:10.1038/s41598-017-02683-3

38. Peg X, Xu E, Liang W, et al. Identification of FAM3D as a new endogenous chemotaxis agonist for the formyl peptide receptors. J Cell Sci. 2016;129(9):1831–1842. doi:10.1242/jcs.183053

