## Supplemental Figure 1 for "Biomarker Identification by Proteomic Analysis of Vitreous Humor and Plasma in Diabetic Retinopathy"

**Supplementary online material: Biomarker Identification by Proteomic Analysis of Vitreous Humor and Plasma in Diabetic Retinopathy**

Supplementary eTables: Refer to eSummplentary Excel file

### Supplementary eFigure 1: Sample Demographics

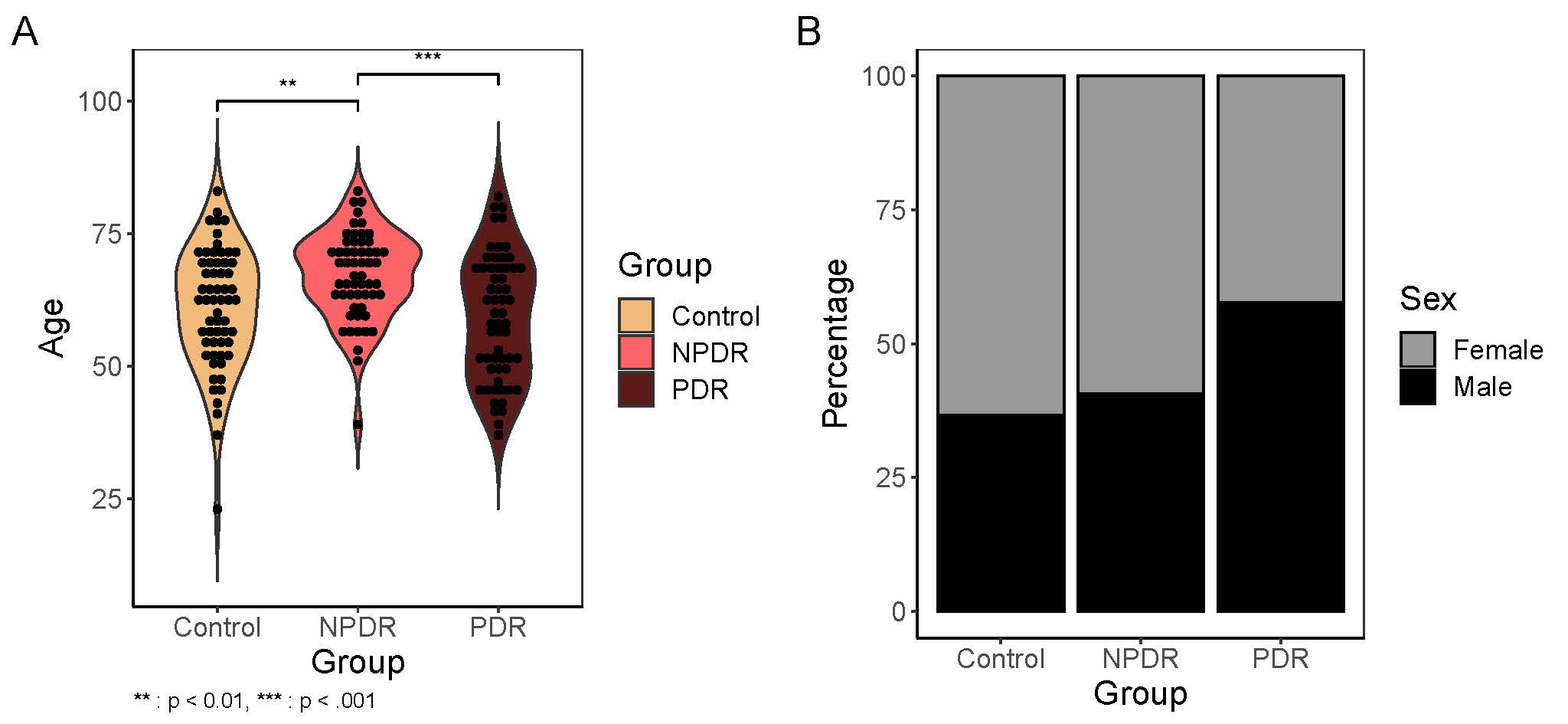

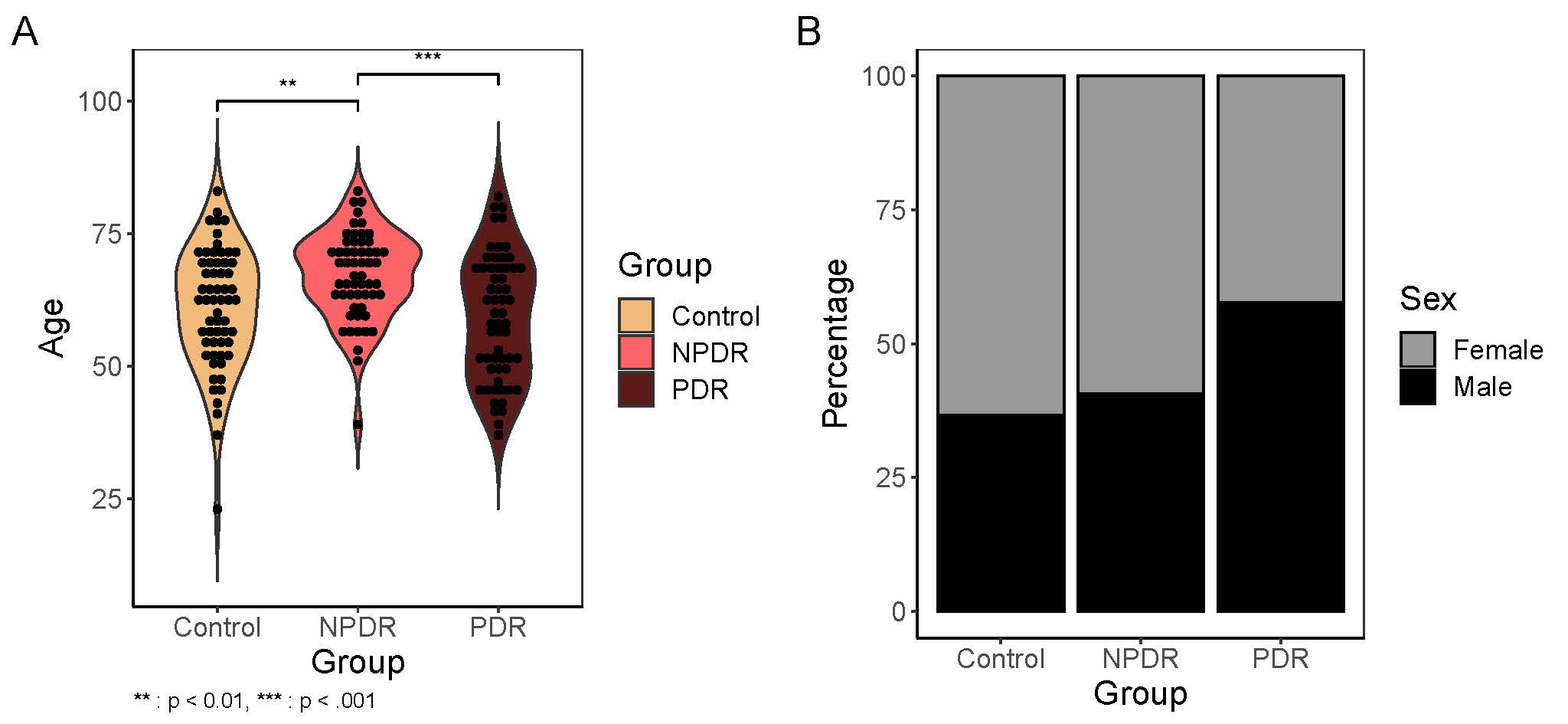

The age (Supplementary eFigure 1A) and gender (Supplementary eFigure1B) distribution for each group. Note: ** indicates p<0.01, *** indicates p<0.001.NPDR, nonproliferative diabetic retinopathy; PDR, proliferative diabetic retinopathy.
